# Dynamic involvement of premotor and supplementary motor areas in bimanual pinch force control

**DOI:** 10.1101/2022.11.14.516422

**Authors:** Anke Ninija Karabanov, Gaetana Chillemi, Kristoffer Hougaard Madsen, Hartwig Roman Siebner

## Abstract

Many bimanual activities of daily living require quick shifts between symmetric and asymmetric motor output generated by the right and left hand. Bimanual motor control has been mostly studied during continuous repetitive tasks, while little research has been carried out in experimental settings requiring dynamic changes in motor output generated by both hands. Here, we performed functional magnetic resonance imaging (MRI) while healthy volunteers performed a visually guided, bimanual pinch force task. This enabled us to map functional activity and connectivity of premotor and motor areas during bimanual pinch force control in different task contexts, requiring mirror-symmetric or inverse-asymmetric changes in discrete pinch force exerted with the right and left hand. The bilateral dorsal premotor cortex showed increased activity and effective coupling to the ipsilateral supplementary motor area (SMA) in the inverse-asymmetric context compared to the mirror-symmetric context of bimanual pitch force control. Compared to unilateral pinches, all bimanual pinch force conditions were characterized by an increased level of activation in the bilateral primary motor hand area (M1-HAND) and stronger coupling from SMA to the ipsilateral M1-HAND. Task-related activity of a cluster in the left caudal SMA scaled positively with the degree of synchronous initiation of bilateral pinch force adjustments, irrespectively of the task context. The results suggest that the dorsal premotor cortex mediates increasing complexity of bimanual coordination by increasing coupling to the SMA-M1 network.

## Introduction

The ability to interact with objects is at the core of most human activities. Dexterous object manipulation, like tying a bow or opening a jar, requires coordination of both hands, often in a complex mixture of symmetric and asymmetric hand movements, that are performed towards a common goal. However, bimanual tasks resembling natural-object oriented actions are underrepresented in bimanual research (Maes et al., 2017). Existing bimanual studies traditionally focus on continuous movements that allow to contrast temporal coordination constraints (e.g. in-phase versus anti-phase finger tapping) or spatial coordination constraints (e.g. circle rotation movements) during steady periods of symmetric or asymmetric coordination (e.g. (Kelso, 1984, Ullen and Bengtsson, 2003, Bengtsson et al., 2004) for review see: (Swinnen and Gooijers, 2015, Rueda-Delgado et al., 2014, Swinnen and Wenderoth, 2004). While these tasks allow robust comparisons between different coordinative patterns, they may underestimate the challenge that quickly shifting coordination patterns poses for the motor system.

Cyclic, continuous bimanual movements result in a stronger activation of the supplementary motor area (SMA) and cingulate motor area (CMA), dorsal premotor cortex (PMd) and motor cerebellum compared to unimanual movements, and the same regions also show stronger activity when contrasting asynchronous bimanual movements with synchronous ones (Sadato et al., 1997, Goerres et al., 1998, Jancke et al., 2000, Ullen et al., 2003). Functional connectivity between these regions is also modulated during bimanual motor control: Symmetric bimanual tapping leads to stronger intra- and interhemispheric coupling between SMA, primary motor cortex (M1) and PMd (Grefkes et al., 2008) than unimanual tapping. Connectivity studies comparing synchronous to asynchronous bimanual tapping report an increased connectivity across the midline during asynchronous movements (Serrien, 2008). EEG data suggests that additional neural resources are recruited at transition points when shifting between synchronous and asynchronous coordination patterns (Banerjee et al., 2012, Aramaki et al., 2006). These findings imply that dynamic bimanual paradigms that incorporate frequent transitions can improve the current understanding of the cortical networks supporting dynamic bimanual control and dexterous object manipulation.

Premotor areas, like SMA and PMd, are not only implicated in bimanual control but are involved in the planning of manual movements in general (Halsband and Lange, 2006): SMA is activated strongly during internally guided movements, whereas PMd is more implicated during tasks with a focus on external, visual cues (Wymbs and Grafton, 2013). In terms of bimanual motor control, this implies that a research focus on movements that rely strongly on internal synchronization (e.g. tapping or circle drawing) may show a different pattern of PMd-SMA interaction (Bengtsson et al., 2004, Meister et al., 2010) (Debaere et al., 2003, Wenderoth et al., 2004, Diedrichsen et al., 2006) than bimanual tasks that resemble natural object manipulation and therefore rely stronger on visual cues (Karabanov et al., 2019) (Theorin and Johansson, 2007, Theorin and Johansson, 2010).

Little is known about how connectivity patterns between PMd and SMA are modulated during visually guided bimanual movements. Especially differences in connectivity patterns between the left and the right hemisphere are of interest as several studies suggest a hemispheric asymmetry for bimanual control (Rueda-Delgado et al., 2014; Serrien, Ivry, & Swinnen, 2006). During symmetric bimanual reaching movements, the left hemisphere seems to have a leading role, at least in right handers, and bilateral reaching movements are encoded stronger in the left hemisphere (Blinch, Flindall, Smaga, Jung, & Gonzalez, 2019; Merrick et al., 2022). Better bimanual coordination is also associated with faster signal transmission from the left M1 to the right hemisphere but not vice versa (Bortoletto et al., 2021). On the other hand, secondary motor areas in the right hemisphere seem to play a crucial role during asymmetric bimanual coordination. Transcranial magnetic stimulation (TMS) induced disruption of the right PMd caused more interference in asymmetric bimanual movement coordination than TMS over the left PMd in both left and right handers (van den Berg, Swinnen, & Wenderoth, 2010). The right hemisphere also displays dominance for the executive control of attention (Spagna, Kim, Wu, & Fan, 2020) which may impact how visual feedback information about the accuracy of the bimanual movements is processed.

To investigate the effect of dynamically changing bimanual coordination requirements on regional activity and interregional connectivity in visuo-motor networks, we developed a novel, a visually guided, dynamic pinch force task that allows to test a wide range of bimanual coordination contexts (mirror-symmetric, inverse-asymmetric and unimanual movements, with one prime mover). We asked healthy volunteers to perform the visually guided, bimanual pinch force task during whole-brain fMRI. Using mass-univariate analysis and dynamic causal modeling (DCM), we tested the hypothesis that complex bimanual movements, during which the right and left hands have to exert different levels of pinch force coordination (e.g. inverse-asymmetric movements), result in a PMd-centered network activation. Additionally, we wanted to explore if changes in connectivity patterns are driven by the dominant hemisphere.

## Methods

### Participants

Thirty-one participants were recruited for the study. Six participants were excluded due to technical issues leading to incomplete data sets. Hence the final sample contained 25 participants (10 women; mean age: 26.1 ± 6.1std). Participants were recruited via an ad on a homepage for recruitment of research participants (www.forsoegsperson.dk). All participants were right-handed, between 18-40 years of age, had no personal history of diagnosed neuropsychiatric disorders, and were not taking neuroactive medication. Participants received detailed information and instructions about the experiment, signed informed consent and were reimbursed for their participation. The study was approved by the Regional Committee on Health Research Ethics of the Capital Region in Denmark (H-16025401).

### Bimanual Pinch Grip Task

To assess dynamically changing coordination requirements, participants performed a visually cued bimanual pinch-grip task. Participants were holding a small, round force sensor (25.4 mm in diameter) in each hand, using a precision pinch-grip with index and thumb. By increasing or decreasing pinch force in each hand participants could control the size of two visually presented semi-circles on a screen, the right semi-circle was controlled by the righthand and the left semi-circle by left. The raw sampling rate of the force sensor was 1024 Hz but the visual signal displayed on the screen was an average over the last 100 samples (e.g. around 0.1 seconds), this smoothing of the force data was induced to minimize electrical background noise. The experimental task required matching the two force-controlled lines to two target semi-circles. The visual semi-circles were adjacent to each other, so that they formed one large circle if the same fraction of maximal voluntary contraction (MVC) was applied by both hands (Figure 1A). At baseline, the task required the participants to keep an isometric pinch contraction of 4% MVC in both hands to match the targe. An increase in the radius of the target semi-circle indicated that the pinch had to be increased to 6% MVC. Correspondingly, a decrease in the target semi-circle indicated that pinch force had to be decreased to 2% MVC. This resulted in nine different conditions (see Figure 1A): Four bimanual conditions in which both hands changed pinch force (Bimanual Symmetry: force increase and force decrease; Bimanual Inverse-Asymmetry: right hand increase (left hand decrease) and vice versa), four unimanual conditions in which only one hand changed pinch force, while the other one maintained a isometric contraction (Left Hand: force increase and force decrease; Right Hand: force increase and force decrease;) as well as a baseline condition where both hands maintained an isometric force at baseline. Pinch-force for each condition had to be held for 2 seconds and was followed by a return to baseline, also lasting for 2 seconds (± 400 ms) before target semi-circles moved to indicate next position. Preliminary analysis of the fMRI data did not indicate significant differences in brain activity between conditions belonging to the same “bimanual category” but with opposing force direction (e.g. Bimanual Symmetry: Force Increase vs. Bimanual Symmetry: Force decrease) even at very liberal thresholds and hence we combined the nine conditions into five general “movement context” conditions: Bimanual Symmetry, Bimanual Asymmetry, Left Hand, Right Hand and Baseline independent of force direction.

**Figure 1:**
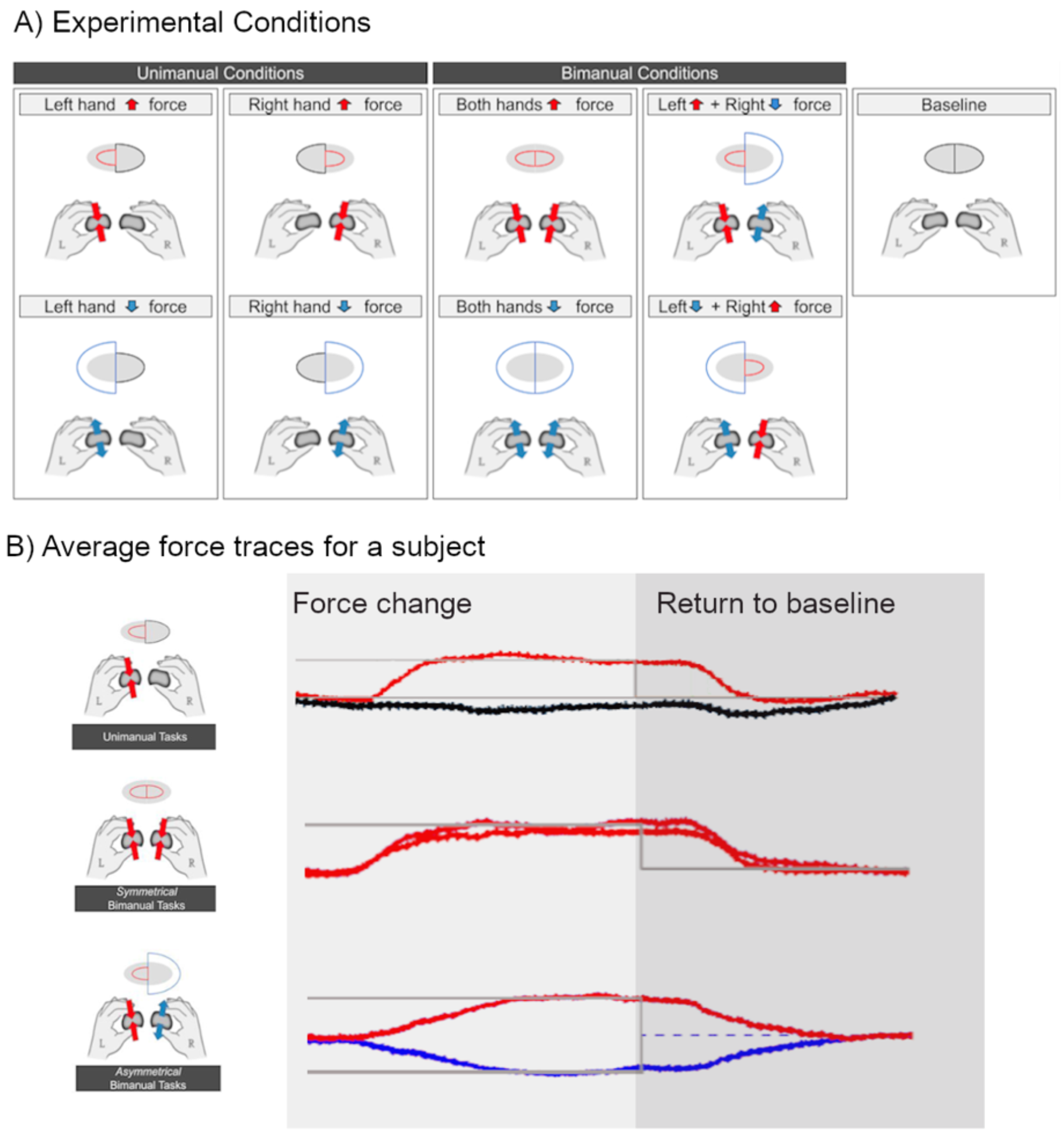
A) Shows the nine different experimental conditions that can further be simplified in Left Hand, Right Hand, Bimanual Symmetry, Bimanual Asymmetry and Baseline. B) Shows an example for the averaged force traces from a representative subject durng Left Hand, Bimanual Symmetry _Force Increase_ and Bimanual asymmetry _RightHand_Force Decrease._

### Experimental procedure

Participants received general information about the experiment and signed the consent form before they were placed in supine position inside the scanner. Participants were given the force sensors and instructed on how to use them while they were lying in the scanner with their arms resting at their sides on supportive cushions. After that, the maximal voluntary contraction (MVC) for the left and the pinch grip was determined using the force sensors. MVC was used as an individual reference for pinch grip strength and the subsequent bimanual pinch force task. After a short familiarization with the task, a structural T1-weighted scan was performed during which the participants were allowed to release the force sensors. After the structural scan participants were instructed to grip both force sensors and underwent three fMRI sessions, separated by brief breaks. Each bimanual pinch force session consisted of five short blocks in each of which the 9 conditions were presented three times in randomized order (110 sec). The short blocks were separated by 30 second pause intervals where participants could release the force sensors to avoid muscle fatigue. Sessions were separated by short breaks of 2-3 minutes where participants could release the force sensors. Across the three sessions participants were in total presented with 45 repetitions of each of the nine conditions (3 per block x 5 blocks x 3 sessions). The scanning session was concluded with three diffusion-weighted scans during which participants could release the force sensors.

### Analysis of behavioral data

The force data for the left and right hands was extracted continuously throughout the experiment. To quantify performance for each condition accuracy (ACC) and reaction time (RT) were determined for each condition and each hand. Accuracy was calculated by averaging the 45 force traces belonging to the same condition in each participant. After that, the area between the target trace and the force trace was calculated for the last 500ms of the trial. The time interval was chosen to ensure that participants had reached a steady force. RT was also calculated using the average individual force trace for each condition. The start of a dynamic movement (RT_start_) was determined as the earliest time-time point after the target force indicated the start of a new movement where the target force deviated more than 15% compared to the previous 100ms. Similarly, return to baseline (RT_end_) was determined as the earliest time point after the target stimulus went back to baseline where the force deviated more than 15% compared to the previous 100 ms. This procedure resulted in one RT start, RT stop, and an ACC score per condition and per hand for each participant. Values more than 2 standard derivations away from the mean were excluded from further analysis (maximally 4.6% of data points per variable).

As one main focus of the study was investigating different bimanual coordination patterns we focused the main behavioral analysis on symmetric and asymmetric bimanual movements but for the interested reader, a complimentary more detailed analysis including all unimanual and bimanual conditions is included in the supplementary material (S1). For each dependent variable (RT_start_, RT_stop_, and Acc) a repeated three-way ANOVA with the independent factors Bimanual Context (Bimanual Symmetric, Bimanual Asymmetric), Hand (Left, Right), and Force (Force Increase, Force Decrease) was calculated to specifically tested the difference between the bimanual conditions. Normality was assessed using the Shapiro-Wilk test. The variables RT_end_ and ACC only passed the Shapiro-Wilk test after log transformation, hence log-transformed values were used in these analyses. A Fisher’s LSD test was used for posthoc testing in case of significant main effects or interaction. All behavioral data were analyzed using the statistical software package R (https://www.r-project.org). The significance threshold for null hypothesis testing was set to p < 0.05.

### Image acquisition

Data acquisition was performed at the Danish Research Centre of Magnetic Resonance, Hvidovre Hospital, Copenhagen, Denmark using a 3T MR scanner (Philips Magnetom Archiva, Best, Netherlands) equipped with a 32-channel array receive coil. Each scan include a structural T1-weighted sequence (MPRAGE; FOV: 245 mm; 245 slices, TR/TE: 5.9/2.7; resolution 0.85×0.85×0.85mm^3^; flip angle: 8 deg.; TI: 747.6 ms), three T2*-weighted echo planar imaging (EPI) sequences utilizing gradient echo (FOV: 192 mm; 42 transverse slices acquired in interleaved order, TR/TE: 2000/30; in-plane resolution:3×3 mm^2^, 3mm slices, no slice gap, flip angle: 90 deg.) and three diffusion-weighted sequences (FOV: 224 mm; 61 directions; TR/TE: 10710/85; resolution: 2.3 × 2.3 × 2.3mm3, flip angle: 8 deg. b = 2000s/mm2 (62 directions), b =1000 s/mm2 (62 directions), b = 300 s/mm2 (6 directions). Results of the diffusion-weighted data will be reported elsewhere.

### Image data analysis

#### Preprocessing

fMRI data were analyzed using Statistical Parametric Mapping (SPM12; Wellcome Trust Center for Neuroimaging, London, UK) implemented in MATLAB R2020a (MathWorks). First, both the MPRAGE and the EPI sequences were manually reoriented according to the location and orientation of the anterior/posterior commissure. After that, all fMRI volumes were slice time corrected to adjust for the sequence of acquisition by sinc interpolating the voxel activation of each slice to the same time point (Henson et al., 1999). The resulting images were realigned, segmented using the SPM12 Tissue Probability Map (Ashburner and Friston, 2005), normalized, and smoothed with an isotropic 6mm full-width half-maximum Gaussian kernel to decrease residual inter-subject differences and to increase the signal-to-noise ratio.

#### Analysis of task-related fMRI data

fMRI data were analyzed using the mass-univariate general linear model (GLM). With the help of default SPM synthetic hemodynamic response function a design matrix was generated. Residual motion was modeled from the rigid body realignment procedure (six parameters) (Friston et al., 1996). Additionally, a high-pass filter modeled low-frequency trends originating from scanner drift. As preliminary analysis did not show significant activations due to force direction (even at a liberal threshold of 0.01 uncorrected) differences in force were collapsed for the different bimanual movement contexts. This resulted in five conditions modeling the bimanual movement context in the final GLM: Bimanual Symmetry, Bimanual Asymmetry, Right Hand, Left Hand, Baseline. To decouple the effects of the Bimanual Coordination Condition, form the error to the target, we included two additional regressors that modeled the continuous error during tracking convolved with the hemodynamic response function for the right and left hand separately. This allowed us to also investigate where BOLD signal variations correlated with the target error. Correlations between the tracking error and BOLD signal can be sound in the supplementary material (S2). For all fMRI data multiple comparisons were corrected for using the family-wise error correction method as implemented in SPM12 (threshold set at p < 0.05).

#### Correlations with behavior

We performed an exploratory analysis to identify brain regions where regional brain activity scales with the temporal synchronization of bimanual pinch force control. We calculated the absolute difference in RT_Start_ between the right and left hand during all bimanual movements (i.e., mirror-symmetric and inverse-asymmetric pinch force adjustments) as an index of bimanual balance of movement onset. Small values indicate synchronous movement initiation, while large values indicate that one hand generally moved faster. This measure of bimanual balance was correlated with task-related activation by bimanual movements, contrasting the individual parameter estimates of bimanual movements and unimanual movements.

For all analyses, we pre-defined PMC, SMA and M1 bilaterally as regions of interest (ROIs), because these regions had been previously identified as core regions of bimanual motor control and learning (Debaere et al., 2004). The ROIs were defined as spheres with a 12 mm diameter centered on the peak stereotactic coordinates of PMd, SMA and M1 reported in (Debaere et al., 2004). For these ROIs, we applied small volume correction using the family-wise error correction method as implemented in SPM12 (threshold set at p < 0.05). The Automated Anatomical Labeling Atlas implemented in the SMP toolbox was used to label active clusters.

#### Dynamic Causal Modeling

Dynamic causal modeling (DCM) was applied to assess changes of connectivity in the bimanual motor network sensitive to changes in bimanual movement context. DCM attempts to model the hidden dynamics underlying changes in fMRI signal based on differential equations, comprising an A-matrix (baseline coupling), a B-matrix (contextual modulation of the connections), and a C-matrix (direct inputs to the system (Friston et al., 2003)). We selected a group of three bilateral ROIs: M1, SMA, PMC as well as a ROI in V5 which served as input region. SMA and PMC were included because interactions between these areas were modulated when comparing simple rhythmic unimanual movements with symmetric bimanual movements (Grefkes et al., 2008). We defined the individual coordinates for each ROIs as the local maximum closest to the coordinates in the previous study. Local maxima were visually inspected according to the same anatomical constraints as listed by Grefkes (Grefkes et al., 2008). The coordinates of the individual ROIs were determined in the respective F-contrast for each subject contrasting Bimanual Symmetric Movements, Bimanual Asymmetric Movements, Left-Hand Movements, and Right-Hand Movements against each other (p < 0.001, uncorrected) (see S3). Time series for the ROIs were extracted using a sphere region (radius = 4 mm) for each subject. The F-contrast showing the common activations for all Bimanual tasks in the ROIs included in the DCM can be found in the supplementary material (S2).

To model bimanual coordination context (Bimanual Symmetric, Bimanual Asymmetric, Left Hand, Right Hand) as a modulatory input to the DCM, fMRI data of all three sessions were concatenated in a new GLM. The model included the same conditions as the first-level GLM described above. Based on previous literature (Grefkes et al., 2008) we constructed four alternative models probing the modulatory effects of Bimanual Asymmetric, Bimanual Symmetric, left Hand, and Right-Hand Movements (for model alternatives see S4). Bayesian model selection (BMS) detected the best model explaining the data (Penny et al., 2004) where differences in free energy (F) indicate evidence for a given model (Friston et al., 2007). After having identified the most likely model, the connectivity parameters (B-matrix) of the winning model were extracted and then entered in a second-level analysis to test if the corresponding coupling parameter was significantly different from each other (Symmetric vs Asymmetric) or from zero. Connections were considered statistically significant if they passed a threshold of p < 0.05 (family wise-error corrected for all connections).

## Results

### Behavioral Performance

The reaction times at the start of the trials indicated significant effects for Bimanual Movement Context, Force Direction and Active Hand: Symmetric bimanual movements were initiated faster than Asymmetric movements (Main effect Bimanual Context: F(1)=33.09; p<0.0001), movements requiring a force decrease were initiated faster than movements requiring a force increase (Main effect Force Direction: F(1)=35.71; p<0.0001) and right handed movements were initiated faster than left-handed movements (F(1)=4.86; p = 0.028). There was a trend for an interaction between Bimanual Context and Force Direction (F(1)=3.77; p=0.053). No other interaction was significant (all p-values > 0.3) (Figure 2A).

**Figure 2:**
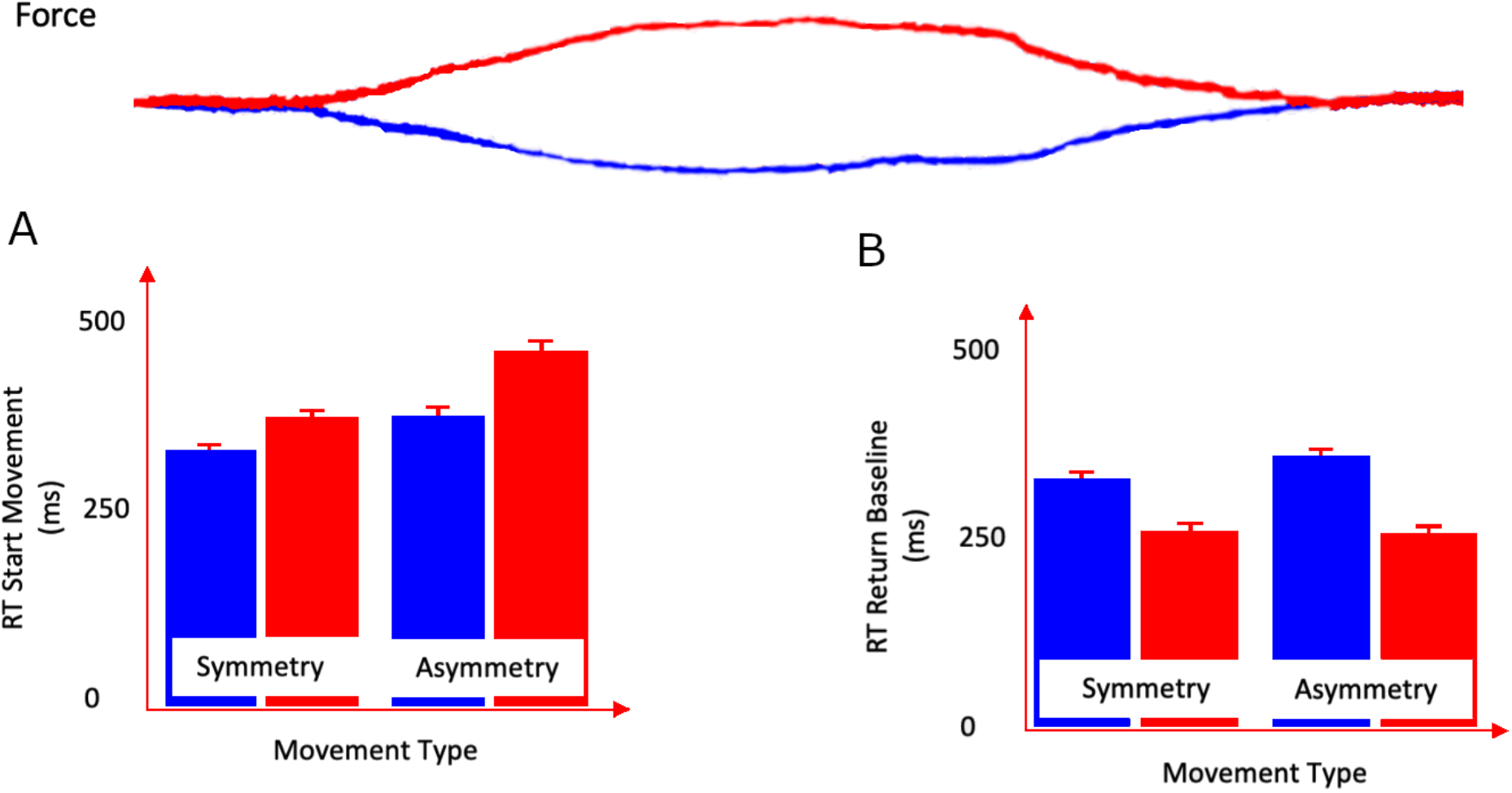
A Shows the average reaction times of an exemplary individual for RT_Start_ (white background) for Bimanual Symmetry and Bimanual Asymmetry separated by force increase (red) and force decrease (blue). B) shows average reaction time for RT_Stop_ (grey background) for Bimanual Symmetry and Asymmetry. Note that the force pattern is reversed for the return to baseline at the end of the trial.

The reaction times for returning to baseline were generally lower than the reaction times at the start of the trial. For force direction they showed a opposite pattern with movements requiring to return to baseline from a force decrease taking longer than movements returning to baseline from a force increase (F(1)=111.9; p>0.0001). Also the interaction between Bimanual Context and Force Direction was significant (F(1)= 4.51; p=0.034). No other main effects of interactions were significant (all p-values > 0.2) (Figure 2B).

Accuracy was higher for movements requiring a force increase than for movements requiring a force decrease (F(1)=101.28; p<0.0001). No other main effects of interactions were significant (all p-values > 0.1). The same pattern also emerged in the analysis that included Unimanual movements as a level in the factor Bimanual Context. This analysis (see S1) showed that reaction times for unimanual movements were generally in between the asymmetric and symmetric conditions (both for RT__start_ and RT__end_). Accuracy was not different between different Bimanual contexts.

### Bimanual Dynamic Movements

To reveal regions activated by bimanual activity irrespective of the specific coordination context voxel-wise activation during all bimanual movements were compared to all unimanual movements. This contrast revealed increased activation in a cluster in the right sensorimotor cortex (SM1; Brodmann Area 3&4 (BA3/4)), a cluster in the occipital cortex, spanning parts of the right primary and secondary visual cortex (V1/V2) and a cluster in the right lateral frontopolar area. An additional cluster in the left SM1 was activated after SVC (Figure 3; Table 1). Bilateral M1 clusters were at least partially driven by a deactivation during unimanual movements of the ipsilateral hand while visual, frontopolar and insular activations were characterized by increased activity during both bimanual movement types. No brain areas were significantly more activated during unimanual than bimanual conditions.

**Table 1:** Bimanual > Unimanual

| Brain Region | Stereotactic coordinates |  |  | Cluster size | Z values |
| --- | --- | --- | --- | --- | --- |
|  | x | y | z |  |  |
| **Primary Sensorimotor Cortex (SM1) R** | 48 | -13 | 56 | 144 | 4.95 |
| **SM1 L** | -39 | -19 | 56 | 49 | 4.14* |
| **Primary/Secondary Visual Cortex (V1/V2) R** | 12 | -82 | 5 | 251 | 4.68 |
| **Lateral Frontopolar Area R** | 33 | 41 | 14 | 68 | 3.93 |

Legend: Z-scores, Cluster size and activation peaks (MNI coordinates) for areas showing higher BOLD response for bimanual movements compared with unimanual movements (Bimanual > Unimanual). $P > 0.05$ FWE-corrected at cluster level. \*Significant at $p > 0.05$ FWE-small volume corrected.

**Figure 3:**
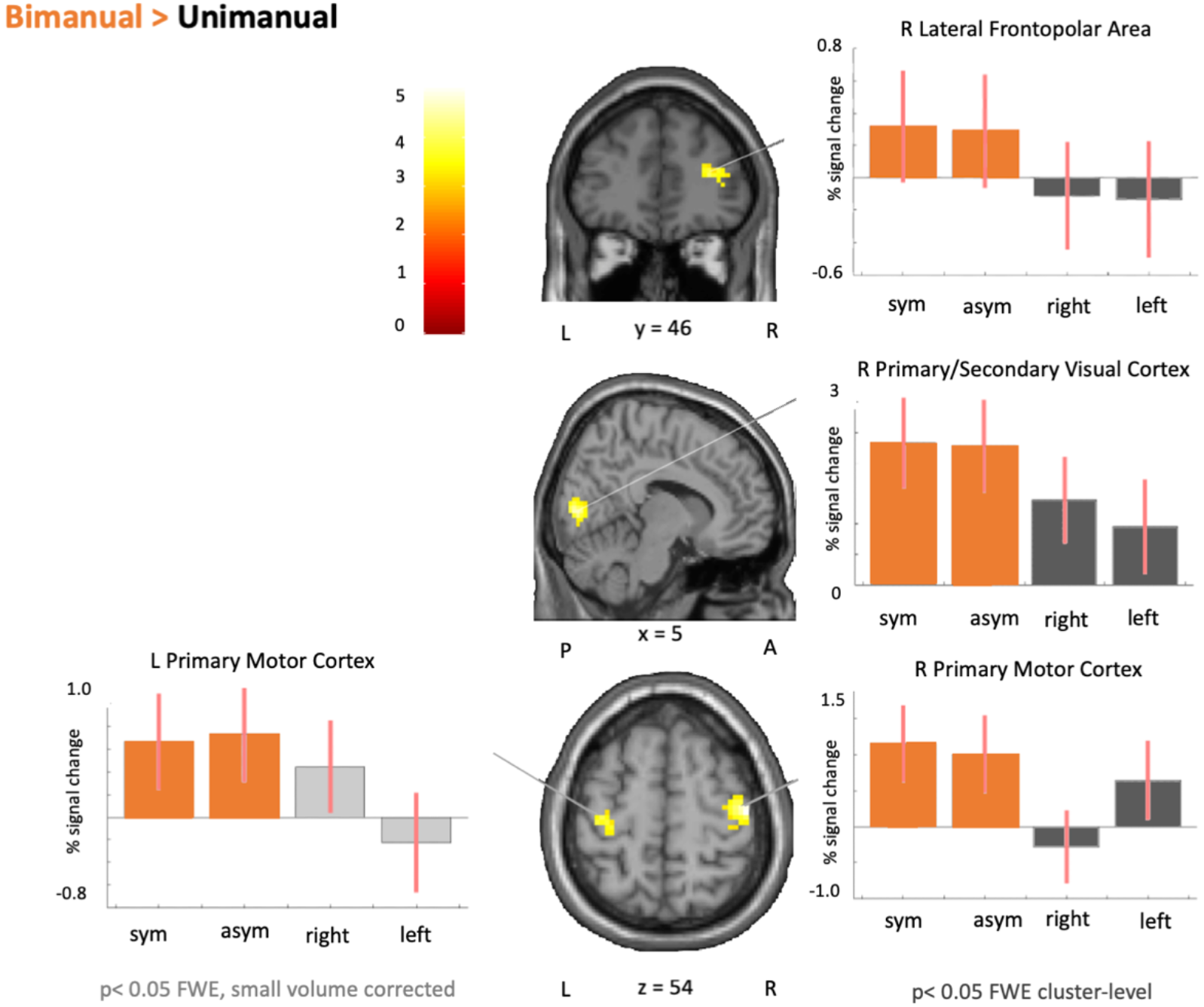
Color-coded statistical parametric maps showing clusters with higher BOLD-signal change for bimanual compared to unimanual movements. The color coding reflects the voxel-specific z-score. The cluster extent of the SPMs are thresholded at 0.05 FWE cluster-level. Bar graphs indicate the relative % signal change in the visualized cluster. Light grey coloring in bar graphs indicate that clusters were only significant at > 0.05 FWE Small-volume corrected. Bimanual regions of interest (ROI) for SVC were taken from Dabaere et al. 2004, Table1. SVC sphere (12mm) around peaks in bimanual coordination network.

### Bimanual Asynchronous Movements

To reveal brain areas activated specifically by bimanual inverse-asymmetric movements, voxel-wise activation patterns compared the bimanual asymmetric condition with the other three conditions. Notably, also this contrast led predominantly to increased activation in hemispheric clusters spanning lower (V1/V2) and higher (V3/4) visual areas. Using the small-volume correction around the predefined ROIs also uncovered bilateral clusters in the dorsal premotor cortex (PMd) (Figure 4 & Table 2). The opposite contrast (comparing mirror-symmetric bimanual with the other conditions) did not reveal any significant activations on a whole brain level or in our predefined regions of interest.

**Table 2:** Asymmetry > Symmetry + Unimanual

| Brain Region | Stereotactic coordinates |  |  | Cluster size | Z values |
| --- | --- | --- | --- | --- | --- |
|  | X | y | z |  |  |
| Fusiform Gyrus /V4 R | 27 | -70 | -4 | 69 | 4.77 |
| V1/V2 L | -21 | -76 | 23 | 92 | 4.17 |
| Area MT/ V5 R | 48 | -73 | 11 | 115 | 4.08 |
| Dorsal Premotor Cortex (dPMC) R | 24 | -7 | 59 | 2 | 3.17* |
| dPMC L | -21 | -4 | 65 | 2 | 3.17* |

**Figure 4:**
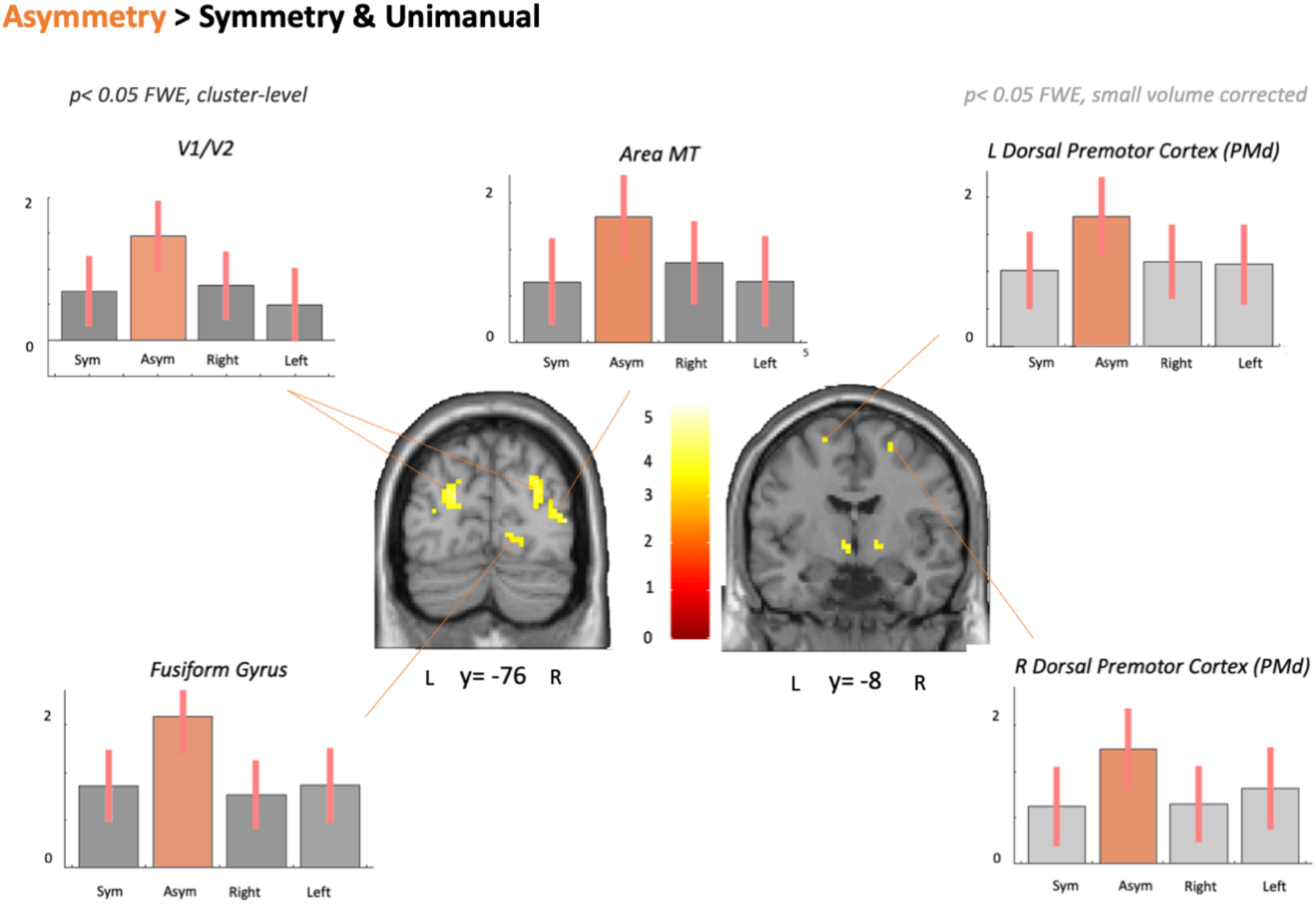
Color-coded statistical parametric maps showing clusters with higher BOLD-signal change for bimanual asymmetric movements compared to the three other conditions. The color coding reflects the voxel-specific z-score. The cluster extent of the SPMs is thresholded at 0.05 FWE cluster-level. Bar graphs indicate the relative % signal change in the visualized clusters. Light grey coloring in bar graphs indicates that clusters were only significant at 0.05 FWE cluster-level after small volume correction (SVC). Bimanual regions of interest (ROI) for SVC were taken from Dabaere et al. 2004, Table1. SVC sphere (12mm) around peaks in bimanual coordination network.

### Correlations with Behavioral Measures

Using regression analysis, we tested which voxel-wise activation patterns were predictive of the degree of bimanual balance. We calculated the absolute ratio in RT_Start_ between mirror-symmetric and inverse-asymmetric bimanual movements to quantify the intermanual synchronization of pinch onset. Task related activity in a small cluster in the posterior part of the left caudal SMA showed a positive linear relationship between task related activity and the degree of intermanual synchronization: Individuals, in whom movement onsets of right and left pinch were highly synchronized, showed a stronger engagement of the left posterior SMA than individuals who showed poor intermanual synchronization (r = - 0.61, p < 0.05, FWE, small volume corrected for ROI; Cluster: x= -9; y= -19, z = -62 and z = 4.47; cluster size 113; Fig. 5a). One participant was removed due to an exceptionally high intermanual synchronization index. The correlation remained significant, when the outlier was included in the analysis.

**Figure 5:**
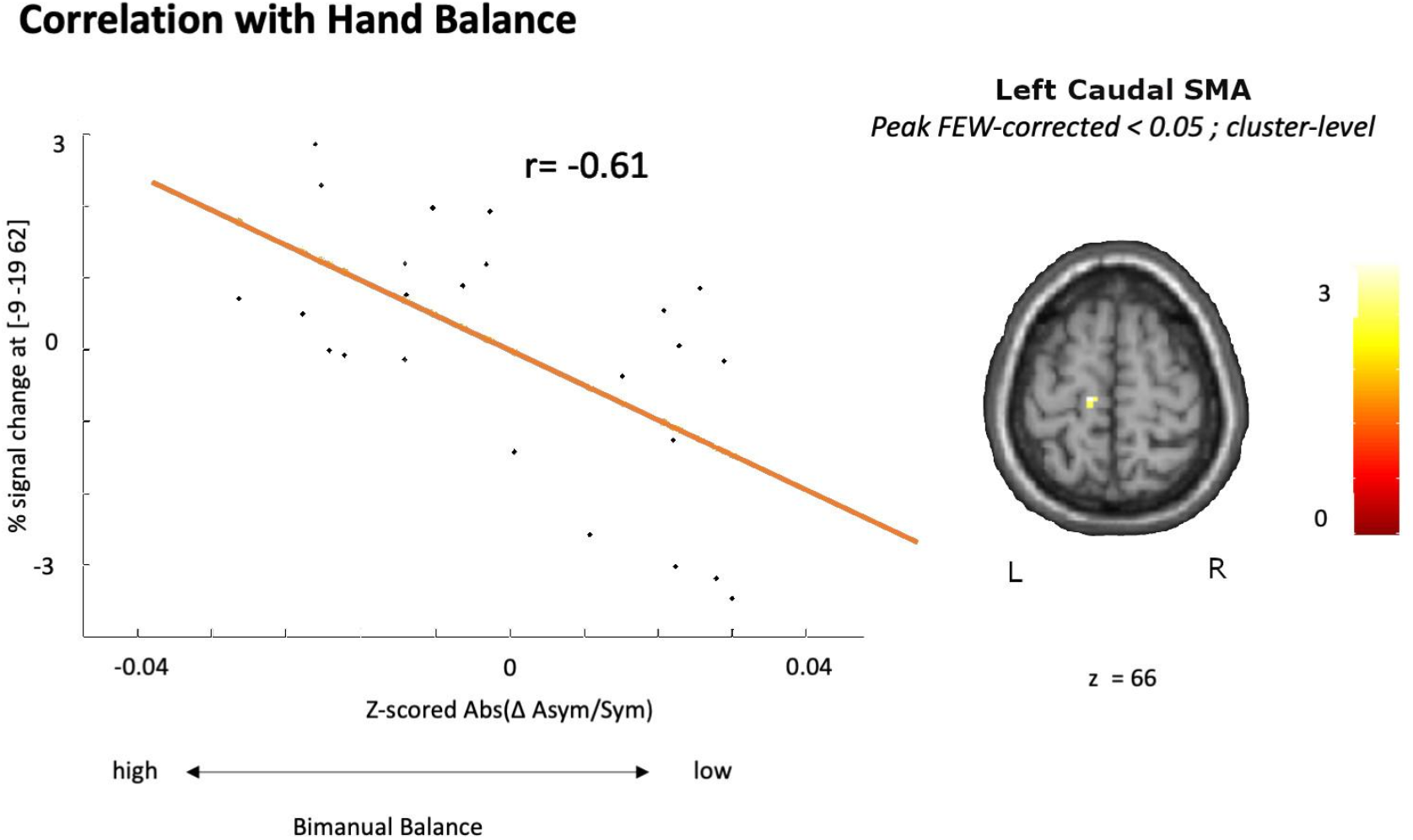
Cluster in left caudal SMA showing a negative linear relationship between hand balance for movement initiation and % signal change. Hand balance was calculated as the absolute differences between RT_onset_ for left and right hand in all bimanual movements. Small volume corrected FWE at p< 0.05)

### Bimanual Context dependent change in effective connectivity

To test the effect bimanual movement context had on network connectivity, DCM analysis were calculated based on previous literature (Grefkes et al., 2008). The winning model had a posterior probability >99.5% and a Bayes Factor above 10. This model assumed a modulatory effect of bimanual movement context on intrahemispheric connections between PMC, SMA and M1 as well as interhemispheric SMA–SMA and M1–M1 connections across hemispheres for all conditions. For unimanual movements, the model additionally assumed a modulatory influence of the contralateral SMA on ipsilateral M1. For symmetric and asymmetric bimanual movements, the model assumed bilateral SMA-M1 connection (See Figure 6 for winning model, for alternative models see supplementary data).

**Figure 6:**
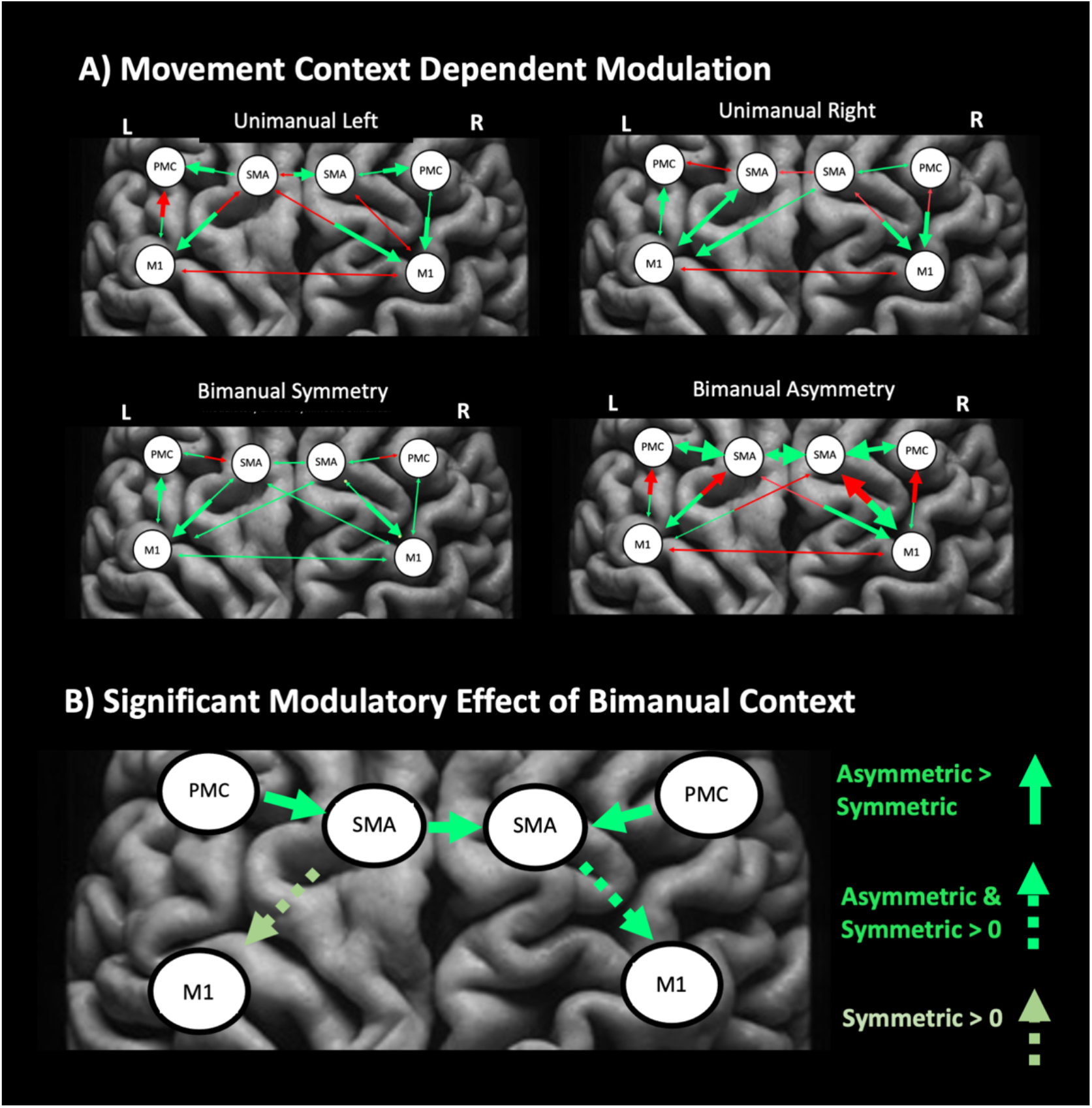
Effective connectivity analysis using DCM. A) Shows the modulatory effect of a task on the connections within each condition. Red arrows indicate negative and green arrows positive modulation. The thickness of the arrow indicates the strength of effective connectivity B) Shows the connections in which the modulatory effect of asymmetric coordination is greater than for symmetric coordination (solid green) as well as connections that are significantly different from zero in either both or one of the bimanual conditions.

Coupling between secondary motor areas was significantly modulated by bimanual movement context. Specifically coupling from the left to the right SMA was upregulated during inverse-asymmetric compared to mirror-symmetric bimanual coordination (p<0.05 FWE). Also coupling from PMd to SMA was upregulated during asymmetric bimanual coordination (p<0.05 FWE) in both hemispheres. Coupling from right SMA to the ipsilateral M1 was not different between symmetric and asymmetric bimanual coordination but was significantly greater than zero in both conditions. The tendency was the same for the left SMA-M1 connection but here, only symmetric coordination increased coupling significantly (p<0.05 FWE). For unimanual coordination, no modulatory effect was significantly different from zero. As modulatory effects were modeled differently for left and right unimanual movements a direct comparison between conditions was not possible. All modulatory weights can be found in S5.

## Discussion

Using a novel visually guided pinch-force task, our results shed new light on the role of premotor areas in visuo-motor coordination of discrete bimanual movements: Trials which required opposite changes in pinch force in the right and left hand engaged the PMd in both hemispheres along with interhemispheric increases in effective connectivity between PMd and SMA. Additionally, interhemispheric coupling from the dominant, left SMA to the non-dominant, right SMA increased during inverse-asymmetric bimanual pinch force adjustments, but without a significant increase in task-related local BOLD activity. The activation of the caudal part of the left SMA in the dominant hemisphere during bimanual movements scaled positively with synchronous initiation of bimanual movements, indicating a general role of this region in the initiation of all types of bimanual movements. Together, our findings support the idea of differential roles of mesial and lateral premotor areas during visually guided, bimanual coordination: While the SMA supports the synchronous initiation of discrete bimanual movements in general, the engagement of the PMd and its influence on ipsilateral SMA increases with complexity, for instance when bimanual force controls requires asymmetric changes in force output.

### Premotor coordination of discrete bimanual pinching movements

Trials requiring a bimanual modulation of pinch force in opposite directions resulted in larger changes in local activity and motor network connectivity than trials requiring discrete symmetric increases or decreases in pinch force. The stronger functional engagement can be attributed to the higher demands on visuomotor mapping which was also evidenced by longer reaction times for asymmetric pinch force adjustments. Our results confirm and extend the results of previous neuroimaging studies that have found increased connectivity between premotor and motor areas when increasing coordinative task difficulty during continuous bimanual tasks (Kiyama et al., 2014, Meister et al., 2010, Rueda-Delgado et al., 2017). Using Structural Equation Modelling (SEM) of fMRI data acquired during visually-paced in-phase and anti-phase tapping, Kiyama et al. (2014) reported increased ipsilateral coupling between PMd and M1 in the left hemisphere in the anti-phase condition. Phase-based connectivity analysis of EEG data revealed increased right hemispheric PMd-M1 coupling when contrasting non-isofrequency and isofrequency dial rotation (Rueda-Delgado et al., 2017). Inter-hemispheric PMd-M1 connectivity has also been shown to be functionally relevant for asymmetric bimanual movements in a study using dual-site transcranial magnetic stimulation (TMS) (Liuzzi et al., 2011): Inter-hemispheric PMd-M1 facilitation of corticospinal excitability, as measured by TMS, was correlated to performance during asymmetric tapping, while inter-hemispheric M1-M1 coupling predicted performance during symmetric and unimanual movements. A notable difference between our results and previously reported results, is that the change in PMd coupling in previous studies was often directed to primary motor areas, whereas we found changes in functional connectivity between premotor areas. The more dynamic and discrete nature of the bimanual pinch force task and the strong reliance on the visual input in each trial increased the functional engagement of premotor areas and thereby the relevance for crosstalk between premotor cortical areas. We found increased left-to-right interhemispheric SMA coupling during inverse-asymmetric bimanual adjustments of pinch force. This finding points to an important role of the SMA in bimanual control and is in good agreement with a previous fMRI study showing stronger coupling from left to right SMA during the preparation of bimanual movements (Welniarz et al., 2019). Clinical studies provide additional strong evidence for a key role of the SMA: Individuals with congenital mirror movements have problems performing discreet asymmetric hand movements and exhibit a lack of left-to-right SMA coupling during bimanual movement execution (Gallea et al., 2013). A focal lesion of the SMA specifically affects the initiation of bimanual movements (Kazennikov et al., 1998, Potgieser et al., 2014). These clinical observations, linking the SMA to the initiation and synchronization of bimanual movements fit well with our finding that the activation level of the caudal left SMA during bimanual pinching is higher, the more the onsets of pinch force adjustments are synchronized between the right and left hand.

### Difference in activity and connectivity between bimanual and unimanual movements

All bimanual pinch movements were characterized by an increased activation of bilateral primary motor hand areas (M1-HAND). During unimanual movements, the M1-HAND ipsilateral to the moving hand showed clear deactivation compared to other conditions, but also the M1-HAND contralateral to the moving hand showed less activation than during bimanual movements. Higher M1-HAND activation in bimanual contexts suggests a context dependent dynamic modulation of interhemispheric inhibition (Carson, 2020) with less inhibition during bimanual compared to unimanual force control. Such a dynamic modulation is functionally relevant as previous studies have shown that reciprocal interhemispheric inhibition is important during unimanual movements (preventing mirror movements) but functionally decremental to bimanual movements (Fling & Seidler, 2012). In our study, activation levels during bimanual pinch adjustments were similar for asymmetric and symmetric conditions indicating that dynamic modulations of BOLD signal in M1-HAND depend more on the general movement context (bimanual vs unimanual) rather than reflecting the specific pattern of force output. This fits well with previous TMS studies that also suggest that interhemispheric inhibition is modulated during bimanual movements but does not differ based on hand symmetry during bimanual tasks (Jordan, Schrafl-Altermatt, Byblow, & Stinear, 2021).

In contrast to other studies, we did not find general activity increases in premotor areas during bimanual movements as reported in many of the early studies (Sadato et al., 1997, Goerres et al., 1998, Jancke et al., 2000, Ullen et al., 2003). This can be attributed to the relatively high complexity of the “unimanual” trials in our task, which required participants to maintain a steady tonic contraction with the non-moving hand. The largest differences between bimanual and unimanual movements in local activity patterns were seen in frontal and occipital regions demonstrating that extrinsic visual coordinates are an integral part in the neural processing required for visually guided bimanual coordination. In fact, visual error feedback can bias participants towards representing bimanual movement patterns, not as effector-dependent muscle coordinates but as extrinsic visual representations of the effector-independent movement outcome (Sakurada and Kansaku, 2021). Since effector-independent coding of movements according to extrinsic coordinates strongly engages the visual cortices (Haar et al., 2017), the strong visual activation clusters may be indicative of the coding of bimanual movement in extrinsic space.

Several studies have compared network coupling between unimanual and (symmetric) bimanual movements (Grefkes et al., 2008, Walsh et al., 2008, Heitger et al., 2013, Maki et al., 2008). Specifically, there is previous DCM work demonstrating that unimanual movements are characterized by increased coupling in pathways contralateral to the moving hand, while the coupling in ipsilateral pathways and between hemispheres is reduced. In contrast, bimanual symmetric movements were characterized by increased interhemispheric coupling between bilateral SMA and M1. The general pattern of positive and negative coupling and the winning models that we observed in healthy individuals performing discrete visually cued pinches with the right and left hand were similar to these previously reported DCM findings, even though not all coupling parameters reached statistical significance in our analysis. Additionally, we found that the interhemispheric coupling between SMA, but not M1, is further upregulated during asymmetric bimanual pinches, requiring an increase in pinch force with one hand and a simultaneous decrease in pinch <u>force</u> with the other hand.

### Hemispheric dominance during bimanual movements

Several studies have suggested that the dominant hemisphere has a leading role during the control of bimanual movements at least for right-handed individual<u>s</u> (Serrien et al., 2003, Walsh et al., 2008, Rueda-Delgado et al., 2014), even though consensus on this is lacking (Rueda-Delgado et al., 2017). Generally, the activity and connectivity changes observed in this study were expressed bilaterally. However, stronger interhemispheric coupling from the dominant SMA to the non-dominant SMA as well as the results implicating the left hemisphere when correlating BOLD activity with accuracy (M1) and synchronicity (paracentral lobule) support the notion that motor areas in the dominant hemisphere have a superior role during the control of bimanual movements. In our study, superiority of the dominant hand during bimanual movements was also indicated by significantly faster reaction times for the right hand independently of the bimanual coordination context. These observations fit well in previous literature (Hoyer and Bastian, 2013, Blinch et al., 2017). Other behavioral studies have also showed that the non-dominant effector reacts more to perturbations of the dominant effector than vice versa (Schaffer and Sainburg, 2021, Yadav and Sainburg, 2014) and that the attention of right handers during bimanual movements is directed to a higher degree towards their dominant right hand ((Buckingham and Carey, 2009, Buckingham et al., 2011).

### Caveats and the effect of force direction

While behavioral data showed that participants were faster to initiate movements involving a force decrease than a force increase, the pinch force direction was not reflected by regional differences in BOLD signal, even at liberal thresholds. Therefore, we pooled data from trials requiring force increase or force decrease in larger categories that investigated bimanual coordination context independent of force direction. When testing a large range of different force levels studies have showed a positive relationship between precision grip force and the BOLD contrast signal, primarily in M1 and S1 (Ehrsson et al., 2001, Sulzer et al., 2011) but the small differences in grip force varying between 2 and 6% of MVF were too weak to induce increase in BOLD contrast, indeed some studies have suggested frontal and parietal areas to be more activated dung very light contractions just above the critical level at which the grasped object would slip (Kuhtz-Buschbeck et al., 2001, Kuhtz-Buschbeck et al., 2008). An additional factor that might have made it difficult to differentiate different force levels was the fixed duration of the 2 seconds for each pinch movement, this made it possible for the participants to anticipate the return to baseline and hence the end of the 2 sec blocks may be “contaminated” by movement preparation for the force-wise mirror opposite return to baseline.

### Conclusions

Using a novel visually guided, bimanual pinch-force task, we demonstrate a bilateral involvement of dorsomedial premotor areas in visuo-motor control of discrete bimanual pinching movements. Enhanced coupling between SMA and M1 appears to be a functional signature shared by all types of bimanual movements. Additional bilateral recruitment of the PMd along with modulations in functional connectivity between PMd and SMA occurs in more demanding bimanual pinching conditions which require an simultaneous increase in pinch force in one hand and a decrease in pinch force in the other hand.

## Acknowledgment

AK holds a 4-year Sapere Aude Starting Grant sponsored by Danmarks Frie Forskningsfond (Grant Nr: 0169-00027B). HS holds a 5-year professorship in precision medicine at the Faculty of Health Sciences and Medicine, University of Copenhagen which is sponsored by the Lundbeck Foundation (Grant Nr. R186-2015-2138). We thank Keenie Ayla Andersen for help with figures and Mikael Novén for helpful discussions of the manuscript.

## Supplementary

### S1: Detailed behavioral model

To compare all three Bimanual Coordination Conditions to each other, each dependent variable (RT start, RT stop, and Acc) was evaluated by a repeated three-way ANOVA with the independent factors Bimanual Context (Bimanual Symmetric, Bimanual Asymmetric, Unimanual), Hand (Left, Right), and Force (Force Increase, Force Decrease). The reaction times at the start of the trials indicated a significant effect for all Bimanual Context and Force Direction: Asymmetric bimanual movements were initiated slower than Symmetric and Unimanual movements (Main effect Bimanual Context: F(1)=12.06; p<0.0004), and movements requiring a force increase were initiated slower than movements requiring a force decrease (Main effect Force Direction: F(1)=41.42; p<0.0001). Finally, there was a trend towards right handed movements being imitated faster than left-handed movements (F(1)=3.30; p = 0.06). No interactions were significant (all p-values > 0.2).

The reaction times for returning to baseline were generally lower than the reaction times at the start of the trial. For force direction they showed a mirror opposite pattern with movements requiring to return to baseline from a force increase taking longer than movements returning to baseline from a force decrease (F(1)=111.9; p>0.0001). Also the interaction between Bimanual Context and Force Direction was significant (F(1)= 4.51; p=0.034). No other main effects of interactions were significant (all p-values > 0.2). Accuracy was also higher for movements requiring a force increase than for movements requiring a force decrease (F(1)=148.2; p<0.0001). No other main effects of interactions were significant (all p-values > 0.2).

### S2: Correlation of task-related activation with Target Error

We wanted to test, in which brain regions the BOLD contrast covaried with the tracking error (i.e., the distance between exerted pinch force and target pinch force) independently of the bimanual context. A single cluster in the left anterior insula scaled its activity with the movement error. The smaller the target error, the higher the activation in the left anterior insula. If the significance level was relaxed (from 0.05 FWE to 0.0001 uncorrected), additional clusters in left PMd (border to frontal eye field), left paramedian upper cerebellum, left lingual and fusiform gyrus and the pars triangularis of the left and right inferior frontal gyrus.

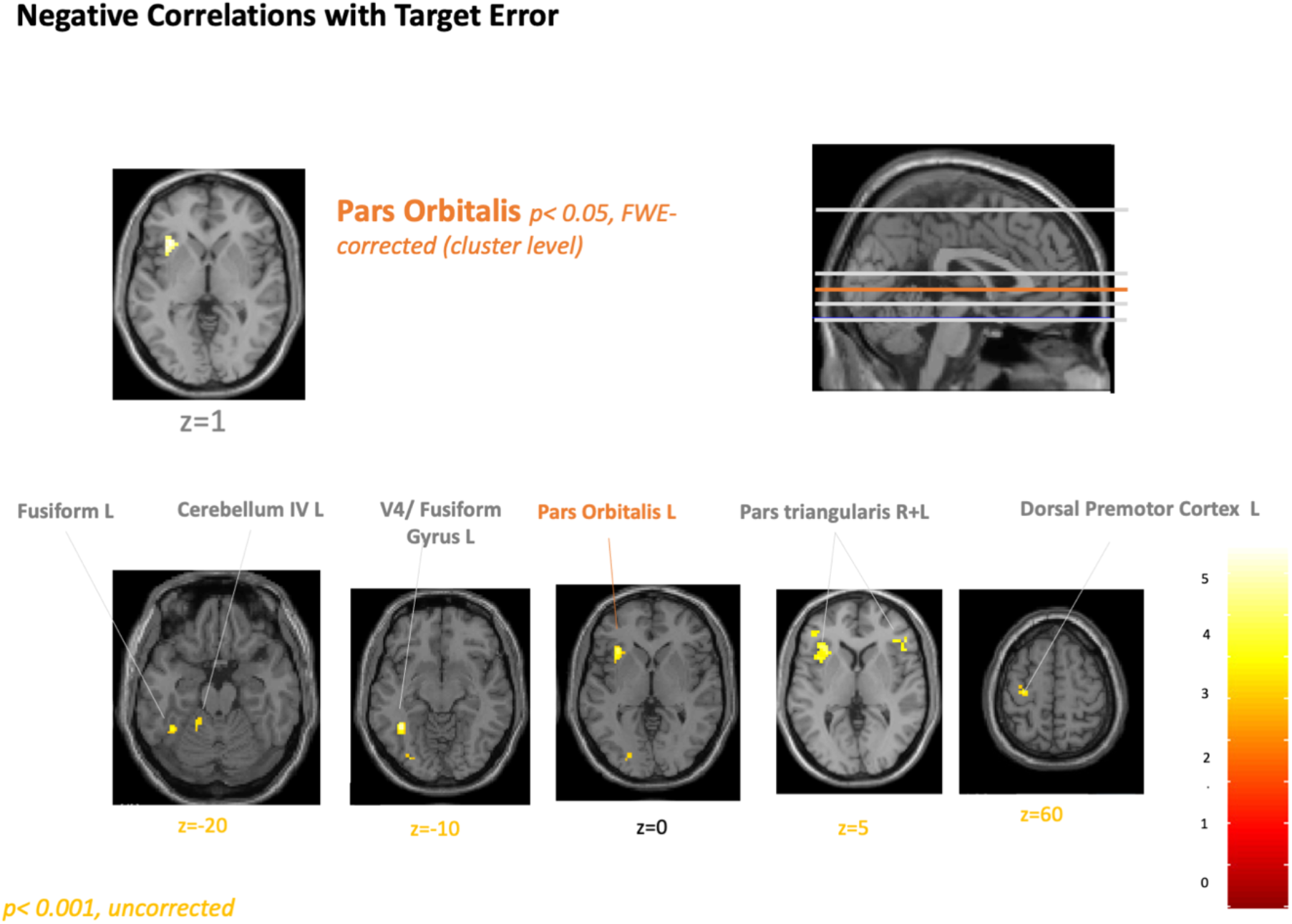

*Clusters showing a negative linear relationship with tracking error (FWE at p< 0*.*05). Clusters labeled in yellow significant at p< 0*.*001 uncorrected*.

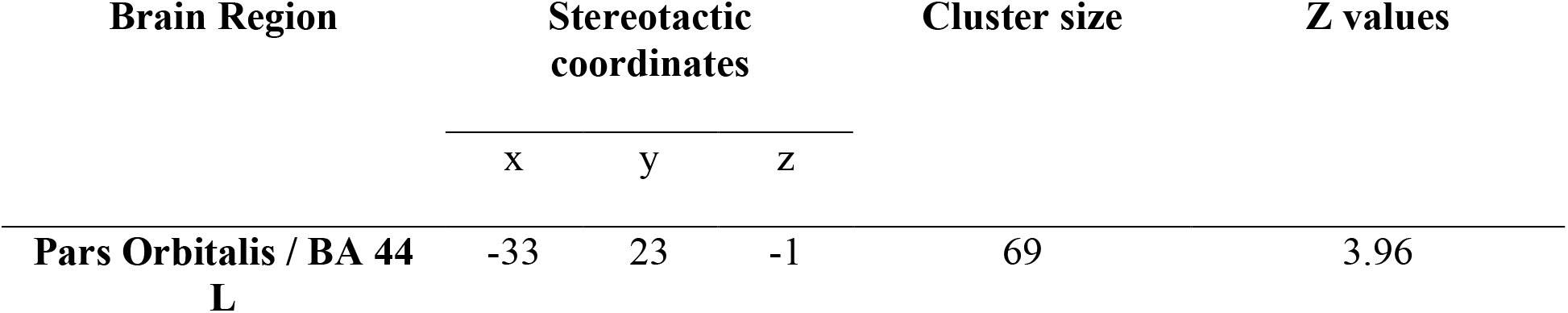
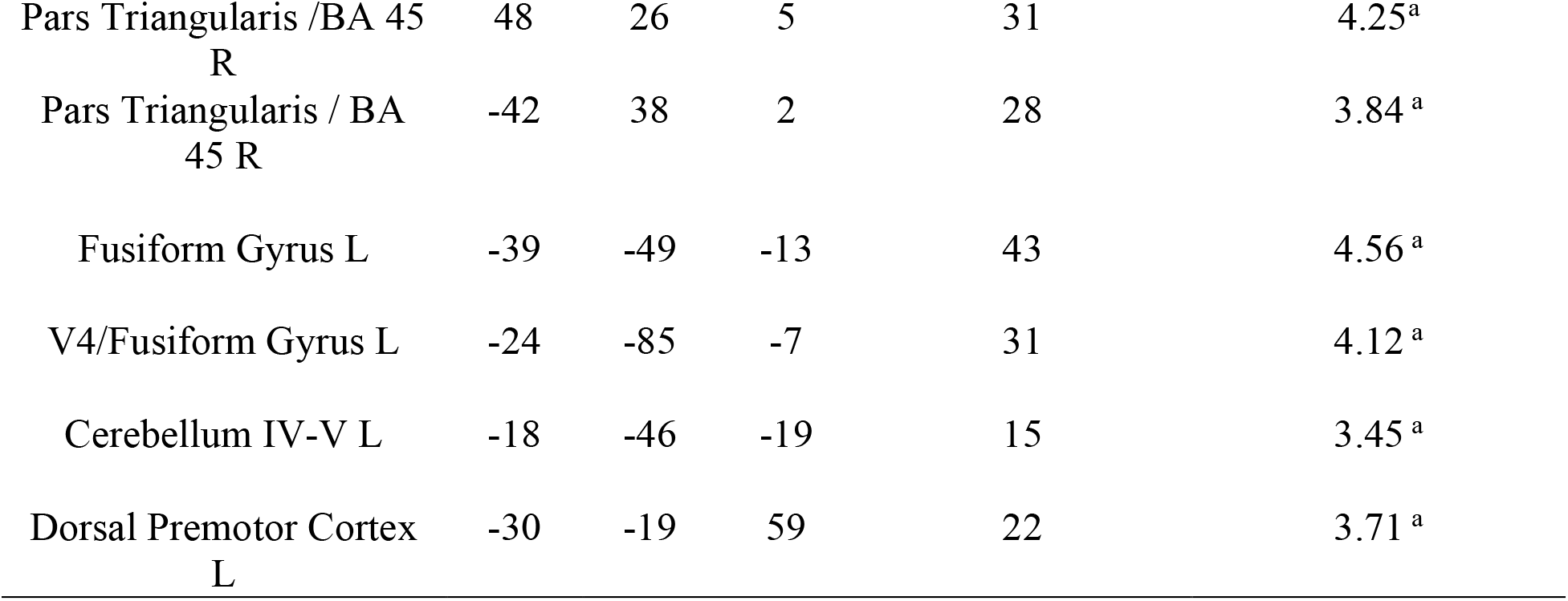

| Brain Region | Stereotactic coordinates |  |  | Cluster size | Z values |
| --- | --- | --- | --- | --- | --- |
|  | x | y | z |  |  |
| **Pars Orbitalis / BA 44 L** | -33 | 23 | -1 | 69 | 3.96 |
| Pars Triangularis /BA 45<br>R | 48 | 26 | 5 | 31 | 4.25<sup>a</sup> |
| Pars Triangularis / BA<br>45 R | -42 | 38 | 2 | 28 | 3.84<sup>a</sup> |
| Fusiform Gyrus L | -39 | -49 | -13 | 43 | 4.56<sup>a</sup> |
| V4/Fusiform Gyrus L | -24 | -85 | -7 | 31 | 4.12<sup>a</sup> |
| Cerebellum IV-V L | -18 | -46 | -19 | 15 | 3.45<sup>a</sup> |
| Dorsal Premotor Cortex<br>L | -30 | -19 | 59 | 22 | 3.71<sup>a</sup> |

### S3: F-contrast between all bimanual contexts

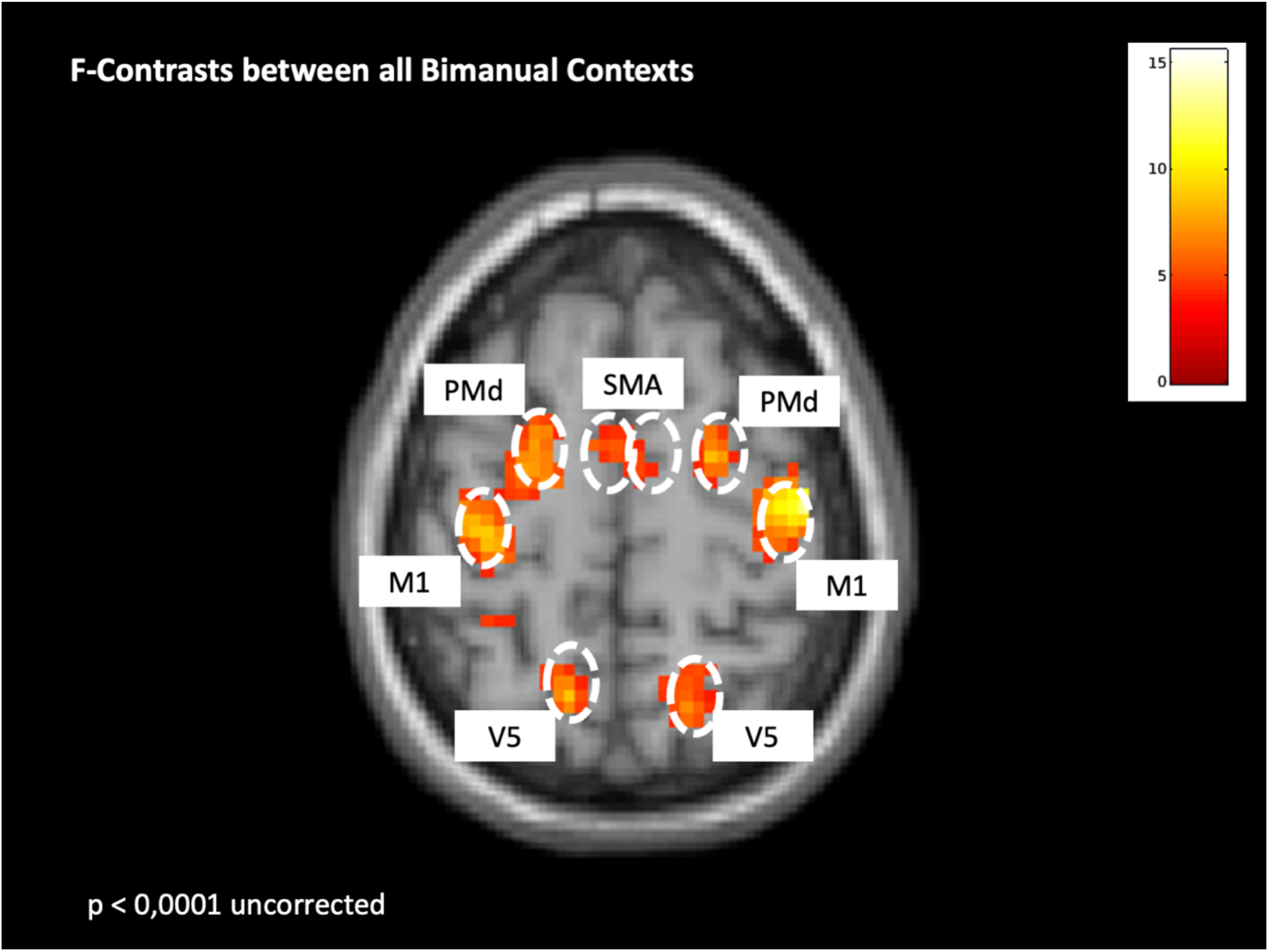

### S4- Alternative DCM models

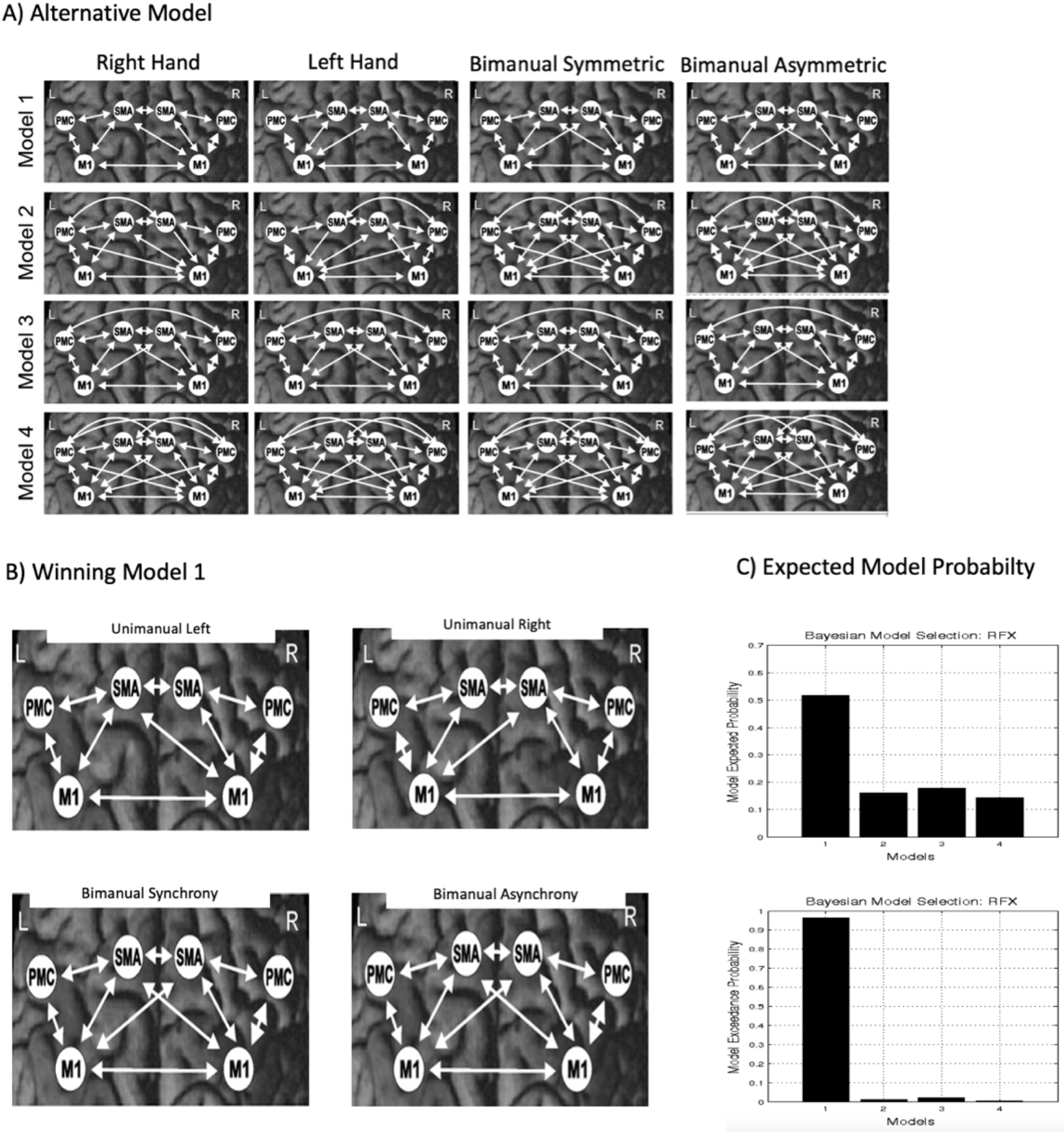

### S5- individual weighting of each connection in the winning model

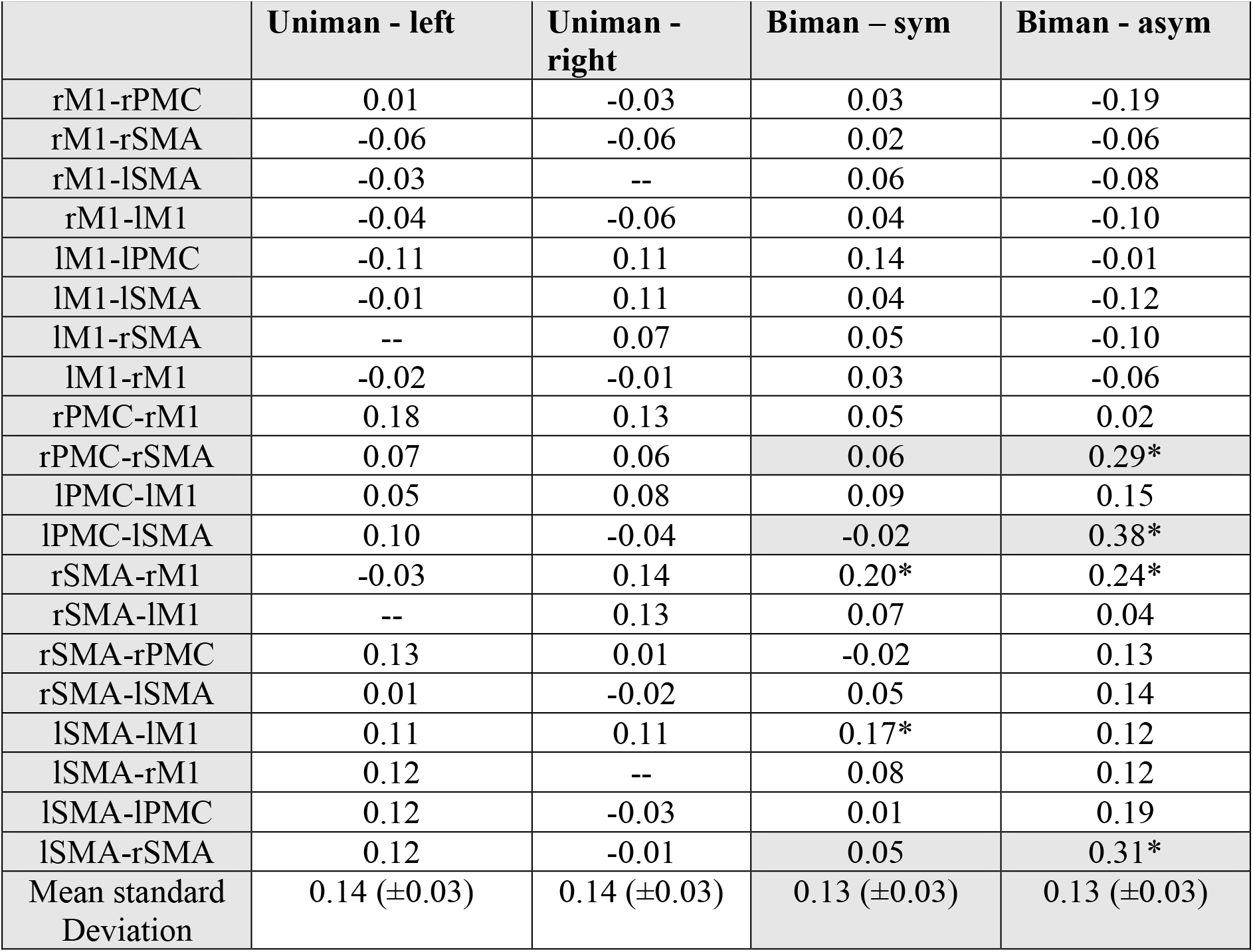

## References

Andersen, K. W. & Siebner, H. 2018. Mapping dexterity and handedness: recent insights and future challenges. Current Opinion in Behavioral Sciences, 20.

Aramaki, Y., Honda, M., Okada, T. & Sadato, N. 2006. Neural correlates of the spontaneous phase transition during bimanual coordination. Cereb Cortex, 16, 1338–48.

Ashburner, J. & Friston, K. J. 2005. Unified segmentation. Neuroimage, 26, 839–51.

Banerjee, A., Tognoli, E., Kelso, J. A. & Jirsa, V. K. 2012. Spatiotemporal re-organization of large-scale neural assemblies underlies bimanual coordination. Neuroimage, 62, 1582–92.

Bastin, J., Deman, P., David, O., Gueguen, M., Benis, D., Minotti, L., Hoffman, D., Combrisson, E., Kujala, J., Perrone-Bertolotti, M., Kahane, P., Lachaux, J. P. & Jerbi, K. 2017. Direct Recordings from Human Anterior Insula Reveal its Leading Role within the Error-Monitoring Network. Cereb Cortex, 27, 1545–1557.

Bengtsson, S. L., Ehrsson, H. H., Forssberg, H. & Ullen, F. 2004. Dissociating brain regions controlling the temporal and ordinal structure of learned movement sequences. Eur J Neurosci, 19, 2591–602.

Blinch, J., De Cellio Martins, G. & Chua, R. 2017. Effects of integrated feedback on discrete bimanual movements in choice reaction time. Exp Brain Res, 235, 247–257.

Blinch, J., Flindall, J. W., Smaga, L., Jung, K., & Gonzales, C. L. (2019). The left cerebral hemisphere may be dominant for the control of bimanual symmetric reach-to-grasp movements. Exp Brain Res, 237(12), 3297–3311. doi:10.1007/s00221-019-05672-2

Bortoletto, M., Bonzano, L., Zazio, A., Ferrari, C., Pedulla, L., Gasparotti, R., Bove, M. (2021). Asymmetric transcallosal conduction delay leads to finer bimanual coordination. Brain Stimul, 14(2), 379–388. doi:10.1016/j.brs.2021.02.002

Buckingham, G. & Carey, D. P. 2009. Rightward biases during bimanual reaching. Exp Brain Res, 194, 197–206.

Buckingham, G., Main, J. C. & Carey, D. P. 2011. Asymmetries in motor attention during a cued bimanual reaching task: left and right handers compared. Cortex, 47, 432–40.

Callaert, D. V., Vercauteren, K., Peeters, R., Tam, F., Graham, S., Swinnen, S. P., Sunaert, S. & Wenderoth, N. 2011. Hemispheric asymmetries of motor versus nonmotor processes during (visuo)motor control. Hum Brain Mapp, 32, 1311–29.

Carson, R. G. (2020). Inter-hemispheric inhibition sculpts the output of neural circuits by co-opting the two cerebral hemispheres. J Physiol, 598(21), 4781–4802. doi:10.1113/JP279793

Debaere, F., Wenderoth, N., Sunaert, S., Van Hecke, P. & Swinnen, S. P. 2003. Internal vs external generation of movements: differential neural pathways involved in bimanual coordination performed in the presence or absence of augmented visual feedback. Neuroimage, 19, 764–76.

Debaere, F., Wenderoth, N., Sunaert, S., Van Hecke, P. & Swinnen, S. P. 2004. Changes in brain activation during the acquisition of a new bimanual coodination task. Neuropsychologia, 42, 855–67.

Diedrichsen, J., Grafton, S., Albert, N., Hazeltine, E. & Ivry, R. B. 2006. Goal-selection and movement-related conflict during bimanual reaching movements. Cereb Cortex, 16, 1729–38.

Ehrsson, H. H., Fagergren, E. & Forssberg, H. 2001. Differential fronto-parietal activation depending on force used in a precision grip task: an fMRI study. J Neurophysiol, 85, 2613–23.

Fling, B. W., & Seidler, R. D. (2012). Task-dependent effects of interhemispheric inhibition on motor control. Behav Brain Res, 226(1), 211–217. doi:10.1016/j.bbr.2011.09.018

Friston, K., Mattout, J., Trujillo-Barreto, N., Ashburner, J. & Penny, W. 2007. Variational free energy and the Laplace approximation. Neuroimage, 34, 220–34.

Friston, K. J., Harrison, L. & Penny, W. 2003. Dynamic causal modelling. Neuroimage, 19, 1273–302.

Friston, K. J., Williams, S., Howard, R., Frackowiak, R. S. & Turner, R. 1996. Movement-related effects in fMRI time-series. Magn Reson Med, 35, 346–55.

Gallea, C., Popa, T., Hubsch, C., Valabregue, R., Brochard, V., Kundu, P., Schmitt, B., Bardinet, E., Bertasi, E., Flamand-Roze, C., Alexandre, N., Delmaire, C., Meneret, A., Depienne, C., Poupon, C., Hertz-Pannier, L., Cincotta, M., Vidailhet, M., Lehericy, S., Meunier, S. & Roze, E. 2013. RAD51 deficiency disrupts the corticospinal lateralization of motor control. Brain, 136, 3333–46.

Goerres, G. W., Samuel, M., Jenkins, I. H. & Brooks, D. J. 1998. Cerebral control of unimanual and bimanual movements: an H2(15)O PET study. Neuroreport, 9, 3631–8.

Grefkes, C., Eickhoff, S. B., Nowak, D. A., Dafotakis, M. & Fink, G. R. 2008. Dynamic intra- and interhemispheric interactions during unilateral and bilateral hand movements assessed with fMRI and DCM. Neuroimage, 41, 1382–94.

Haar, S., Dinstein, I., Shelef, I. & Donchin, O. 2017. Effector-Invariant Movement Encoding in the Human Motor System. J Neurosci, 37, 9054–9063.

Halsband, U. & Lange, R. K. 2006. Motor learning in man: a review of functional and clinical studies. J Physiol Paris, 99, 414–24.

Heitger, M. H., Goble, D. J., Dhollander, T., Dupont, P., Caeyenberghs, K., Leemans, A., Sunaert, S. & Swinnen, S. P. 2013. Bimanual motor coordination in older adults is associated with increased functional brain connectivity--a graph-theoretical analysis. PLoS One, 8, e62133.

Henson, R., Buechel, C., Josephs, O. & Friston, K. 1999. The slice-timing problem in event-related fMRI. NeuroImage 9.

Higo, T., Mars, R. B., Boorman, E. D., Buch, E. R. & Rushworth, M. F. 2011. Distributed and causal influence of frontal operculum in task control. Proc Natl Acad Sci U S A, 108, 4230–5.

Hoyer, E. H. & Bastian, A. J. 2013. The effects of task demands on bimanual skill acquisition. Exp Brain Res, 226, 193–208.

Irmen, F., Karabanov, A., BÖGemann, S., Andersen, E., Madsen, K. H., Bisgaard, T., Dyrby, T. B. & Siebner, H. 2020. Functional and Structural Plasticity Co-express in a Left Premotor Region During Early Bimanual Skill Learning. Frontiers in Human Neuroscience 14:310. doi: 10.3389/fnhum.2020.00310.

Jancke, L., Peters, M., Himmelbach, M., Nosselt, T., Shah, J. & Steinmetz, H. 2000. fMRI study of bimanual coordination. Neuropsychologia, 38, 164–74.

Jordan, H. T., Schrafl-Altermatt, M., Byblow, W. D., & Stinear, C. M. (2021). The modulation of short and long-latency interhemispheric inhibition during bimanually coordinated movements. Exp Brain Res, 239(5), 1507–1516. doi:10.1007/s00221-021-06074-z

Karabanov, A. N., Irmen, F., Madsen, K. H., Haagensen, B. N., Schulze, S., Bisgaard, T. & Siebner, H. R. 2019. Getting to grips with endoscopy - Learning endoscopic surgical skills induces bi-hemispheric plasticity of the grasping network. Neuroimage, 189, 32–44.

Kazennikov, O., Hyland, B., Wicki, U., Perrig, S., Rouiller, E. M. & Wiesendanger, M. 1998. Effects of lesions in the mesial frontal cortex on bimanual co-ordination in monkeys. Neuroscience, 85, 703–16.

Kelso, J. A. 1984. Phase transitions and critical behavior in human bimanual coordination. Am J Physiol, 246, R1000–4.

Kilbreath, S. & Heard, R. 2005. Frequency of hand use in healthy older persons. Australian Journal of Physiotherapy, 51.

Kiyama, S., Kunimi, M., Iidaka, T. & Nakai, T. 2014. Distant functional connectivity for bimanual finger coordination declines with aging: an fMRI and SEM exploration. Front Hum Neurosci, 8, 251.

Kuhtz-Buschbeck, J. P., Ehrsson, H. H. & Forssberg, H. 2001. Human brain activity in the control of fine static precision grip forces: an fMRI study. Eur J Neurosci, 14, 382–90.

Kuhtz-Buschbeck, J. P., Gilster, R., Wolff, S., Ulmer, S., Siebner, H. & Jansen, O. 2008. Brain activity is similar during precision and power gripping with light force: an fMRI study. Neuroimage, 40, 1469–81.

Limanowski, J., Kirilina, E. & Blankenburg, F. 2017. Neuronal correlates of continuous manual tracking under varying visual movement feedback in a virtual reality environment. Neuroimage, 146, 81–89.

Liuzzi, G., Horniss, V., Zimerman, M., Gerloff, C. & Hummel, F. C. 2011. Coordination of uncoupled bimanual movements by strictly timed interhemispheric connectivity. J Neurosci, 31, 9111–7.

Maes, C., Gooijers, J., Orban De Xivry, J. J., Swinnen, S. P. & Boisgontier, M. P. 2017. Two hands, one brain, and aging. Neurosci Biobehav Rev, 75, 234–256.

Maki, Y., Wong, K. F., Sugiura, M., Ozaki, T. & Sadato, N. 2008. Asymmetric control mechanisms of bimanual coordination: an application of directed connectivity analysis to kinematic and functional MRI data. Neuroimage, 42, 1295–304.

Mansouri, F. A., Koechlin, E., Rosa, M. G. P. & Buckley, M. J. 2017. Managing competing goals - a key role for the frontopolar cortex. Nat Rev Neurosci, 18, 645–657.

Meister, I. G., Foltys, H., Gallea, C. & Hallett, M. 2010. How the brain handles temporally uncoupled bimanual movements. Cereb Cortex, 20, 2996–3004.

Merrick, C. M., Dixon, T. C., Breska, A., Lin, J., Chang, E. F., King-Stevens, D., Ivry, R. B. (2022). Left hemisphere dominance for bilateral kinematic encoding in the human brain. Elife, 11. doi:10.7554/eLife.69977

Penny, W. D., Stephan, K. E., Mechelli, A. & Friston, K. J. 2004. Comparing dynamic causal models. Neuroimage, 22, 1157–72.

Potgieser, A. R., De Jong, B. M., Wagemakers, M., Hoving, E. W. & Groen, R. J. 2014. Insights from the supplementary motor area syndrome in balancing movement initiation and inhibition. Front Hum Neurosci, 8, 960.

Rueda-Delgado, L. M., Solesio-Jofre, E., Mantini, D., Dupont, P., Daffertshofer, A. & Swinnen, S. P. 2017. Coordinative task difficulty and behavioural errors are associated with increased long-range beta band synchronization. Neuroimage, 146, 883–893.

Rueda-Delgado, L. M., Solesio-Jofre, E., Serrien, D. J., Mantini, D., Daffertshofer, A. & Swinnen, S. P. 2014. Understanding bimanual coordination across small time scales from an electrophysiological perspective. Neurosci Biobehav Rev, 47, 614–35.

Sadato, N., Yonekura, Y., Waki, A., Yamada, H. & Ishii, Y. 1997. Role of the supplementary motor area and the right premotor cortex in the coordination of bimanual finger movements. J Neurosci, 17, 9667–74.

Sakurada, T. & Kansaku, K. 2021. Attention-dependent switching between intrinsic-muscle and extrinsic-visual coordinates during bimanual movements. Eur J Neurosci, 53, 1922–1937.

Schaffer, J. E. & Sainburg, R. L. 2021. Interlimb Responses to Perturbations of Bilateral Movements are Asymmetric. J Mot Behav, 53, 217–233.

Serrien, D. J. 2008. Coordination constraints during bimanual versus unimanual performance conditions. Neuropsychologia, 46, 419–25.

Serrien, D. J., Cassidy, M. J. & Brown, P. 2003. The importance of the dominant hemisphere in the organization of bimanual movements. Hum Brain Mapp, 18, 296–305.

Serrien, D. J., Ivry, R. B., & Swinnen, S. P. (2006). Dynamics of hemispheric specialization and integration in the context of motor control. Nat Rev Neurosci, 7(2), 160–166. doi:10.1038/nrn1849

Spagna, A., Kim, T. H., Wu, T., & Fan, J. (2020). Right hemisphere superiority for executive control of attention. Cortex, 122, 263–276. doi:10.1016/j.cortex.2018.12.012

Sulzer, J. S., Chib, V. S., Hepp-Reymond, M. C., Kollias, S. & Gassert, R. 2011. BOLD correlations to force in precision grip: an event-related study. Annu Int Conf IEEE Eng Med Biol Soc, 2011, 2342–6.

Swinnen, S. P. & Gooijers, J. 2015. Bimanual Coordination. In: Toga, A. W. (ed.) BRain MApping: An Encyclopedic Refernece London: Academic Press: Elsevier

Swinnen, S. P. & Wenderoth, N. 2004. Two hands, one brain: cognitive neuroscience of bimanual skill. Trends Cogn Sci, 8, 18–25.

Theorin, A. & Johansson, R. S. 2007. Zones of bimanual and unimanual preference within human primary sensorimotor cortex during object manipulation. Neuroimage, 36 Suppl 2, T2–T15.

Theorin, A. & Johansson, R. S. 2010. Selection of prime actor in humans during bimanual object manipulation. J Neurosci, 30, 10448–59.

Tzvi, E., Koeth, F., Karabanov, A. N., Siebner, H. R. & Kramer, U. M. 2020. Cerebellar - Premotor cortex interactions underlying visuomotor adaptation. Neuroimage, 220, 117142.

Ullen, F. & Bengtsson, S. L. 2003. Independent processing of the temporal and ordinal structure of movement sequences. J Neurophysiol, 90, 3725–35.

Ullen, F., Forssberg, H. & Ehrsson, H. H. 2003. Neural networks for the coordination of the hands in time. J Neurophysiol, 89, 1126–35.

Van Den Berg, F. E., Swinnen, S. P., & Wenderoth, N. (2010). Hemispheric asymmetries of the premotor cortex are task specific as revealed by disruptive TMS during bimanual versus unimanual movements. Cereb Cortex, 20(12), 2842–2851. doi:10.1093/cercor/bhq034

Walsh, R. R., Small, S. L., Chen, E. E. & Solodkin, A. 2008. Network activation during bimanual movements in humans. Neuroimage, 43, 540–53.

Welniarz, Q., Gallea, C., Lamy, J. C., Meneret, A., Popa, T., Valabregue, R., Beranger, B., Brochard, V., Flamand-Roze, C., Trouillard, O., Bonnet, C., Bruggemann, N., Bitoun, P., Degos, B., Hubsch, C., Hainque, E., Golmard, J. L., Vidailhet, M., Lehericy, S., Dusart, I., Meunier, S. & Roze, E. 2019. The supplementary motor area modulates interhemispheric interactions during movement preparation. Hum Brain Mapp, 40, 2125–2142.

Wenderoth, N., Debaere, F., Sunaert, S., Van Hecke, P. & Swinnen, S. P. 2004. Parieto-premotor areas mediate directional interference during bimanual movements. Cereb Cortex, 14, 1153–63.

Wymbs, N. F. & Grafton, S. T. 2013. Contributions from the left PMd and the SMA during sequence retrieval as determined by depth of training. Exp Brain Res, 224, 49–58.

Yadav, V. & Sainburg, R. L. 2014. Limb dominance results from asymmetries in predictive and impedance control mechanisms. PLoS One, 9, e93892.

